# PxdA interacts with the DipA phosphatase to regulate endosomal hitchhiking of peroxisomes

**DOI:** 10.1101/2020.02.03.932616

**Authors:** John Salogiannis, Jenna R. Christensen, Adriana Aguilar-Maldonado, Nandini Shukla, Samara L. Reck-Peterson

**Affiliations:** Department of Cellular and Molecular Medicine, University of California San Diego, La Jolla, CA; Howard Hughes Medical Institute, Chevy Chase, MD; The Ohio State Biochemistry Program, The Ohio State University, Columbus, OH; Department of Molecular Genetics, The Ohio State University, Columbus, OH; Section of Molecular Biology, Division of Biological Sciences, University of California San Diego, La Jolla, CA; Section of Cell and Developmental Biology, Division of Biological Sciences, University of California San Diego, La Jolla, CA

**Author notes:** Equal contribution (John Salogiannis, Jenna R. Christensen).

**Keywords:** hitchhiking, endosome, peroxisome, lipid droplet, dynein, kinesin

## Abstract

In canonical microtubule-based transport, adaptor proteins link cargos to the molecular motors dynein and kinesin. Recently, an alternative mode of transport known as ‘hitchhiking’ was discovered, in which a cargo achieves motility by hitching a ride on an already-motile cargo, rather than attaching to a motor protein. Hitchhiking has been best-studied in two filamentous fungi, *Aspergillus nidulans* and *Ustilago maydis*. In *U. maydis*, ribonucleoprotein complexes, peroxisomes, lipid droplets, and endoplasmic reticulum all hitchhike on early endosomes. In *A. nidulans*, peroxisomes hitchhike using a putative molecular linker, PxdA, that associates with early endosomes. However, whether other organelles use PxdA to hitchhike on early endosomes is unclear, as are the molecular mechanisms that regulate hitchhiking in *A. nidulans*. Here we find that the proper distribution of lipid droplets, mitochondria and autophagosomes do not require PxdA, suggesting that PxdA is a molecular linker specific to peroxisomes. We also identify two new *pxdA* alleles, including a point mutation (R2044P) that disrupts PxdA’s ability to associate with early endosomes and reduces peroxisome movement. Finally, we identify a novel regulator of peroxisome hitchhiking, the phosphatase DipA. DipA co-localizes with early endosomes and its early endosome-association relies on PxdA.

## Introduction

The precise spatiotemporal distribution of cargos is critical for cell growth, maturation, and maintenance. Long-distance movement of cargos including vesicles, organelles, mRNAs, and macromolecular complexes is driven by molecular motor-dependent transport on microtubules (Vale, 2003; Hirokawa *et al*., 2009; Cianfrocco *et al*., 2015; Reck-Peterson *et al*., 2018). Microtubules are polarized structures with their “plus” ends located near the cell periphery and “minus” ends embedded near the nucleus at microtubule-organizing centers. In mammalian cells, dozens of kinesin motors carry cargos long distances towards the cell periphery, but a single cytoplasmic-dynein-1 (‘dynein’ here) transports cargos towards the cell center (Vale, 2003; Hirokawa *et al*., 2009; Cianfrocco *et al*., 2015; Reck-Peterson *et al*., 2018).

The filamentous fungus *Aspergillus nidulans* is an ideal model system to study mechanisms of microtubule-based transport (Egan *et al*., 2012a; Peñalva *et al*., 2012). Similar to mammalian cells, but unlike budding yeast, *A. nidulans* uses microtubule-based transport for the distribution and long-distance movement of cargos within its long hyphae. It encodes one dynein, *nudA*, and three cargo-carrying kinesins including a kinesin-1, *kinA* and two kinesin-3’s, *uncA* and *uncB* (Egan *et al*., 2012a; Peñalva *et al*., 2012). Microtubules near the hyphal tips are uniformly polarized with their plus ends oriented outward (Egan *et al*., 2012b), making the directionality of cargo transport and the motors involved easy to identify. We and others have exploited these features by performing forward genetic screens to identify regulators of microtubule-based transport (Morris, 1975; Xiang *et al*., 1994, 1999; Downes *et al*., 2014; Tan *et al*., 2014; Yao *et al*., 2014, 2015; Zhang *et al*., 2014).

The current dogma of microtubule-based transport is that distinct cargos directly recruit molecular motors via adaptors (Fu and Holzbaur, 2014; Reck-Peterson *et al*., 2018; Cross and Dodding, 2019). For example, in mammalian cells, members of the Bicaudal-D and Hook cargo adaptor families link dynein and some kinesins to cargo (Hirokawa *et al*., 2009; Splinter *et al*., 2010; Bielska *et al*., 2014; Hoogenraad and Akhmanova, 2016; Reck-Peterson *et al*., 2018; Cross and Dodding, 2019; Kendrick *et al*., 2019; Siddiqui *et al*., 2019). Altogether, there are dozens of different cargo adaptors that link dynein and kinesin to their cargos. On the other hand, there are relatively few cargo adaptors in filamentous fungi. A single homolog from the Hook family (HookA in *A. nidulans* and Hok1 in *U. maydis*) is the only characterized cargo adaptor for microtubule-based motors in filamentous fungi (Bielska *et al*., 2014; Zhang *et al*., 2014). Hook proteins are one part of the FHF complex (Xu *et al*., 2008; Guo et al., 2016), composed of Fts, Hook, and Fts-Hook-Interacting Protein (FHIP), which links dynein to early endosomes (EEs) via the small GTPase Rab5/RabA in *A. nidulans* (Yao *et al*., 2014). In both *A. nidulans* and *U. maydis*, while EEs are the best-characterized microtubule-based cargo (Wedlich-Söldner *et al*., 2002; Lenz *et al*., 2006; Abenza *et al*., 2009), many other cargos are moved and distributed along the hyphal axis (Egan *et al*., 2012a). How these fungi are capable of properly distributing many cargos despite having few genetically encoded motors and adaptors is unknown. Recently, a non-canonical mechanism of transport termed ‘hitchhiking’ was discovered (Baumann *et al*., 2012, 2014; Higuchi *et al*., 2014; Guimaraes *et al*., 2015; Salogiannis *et al*., 2016). A hitchhiking cargo, rather than connecting directly to an adaptor-motor complex, instead achieves motility by attaching itself to another motile-competent cargo (Salogiannis and Reck-Peterson, 2017). Hitchhiking represents a mechanism that could be employed to move many cargos by a small number of motor-bound cargo in organisms with few genetically-encoded motors and cargo adaptors (Lin *et al*., 2016; Mogre *et al*., 2020).

A number of cargos have been shown to exhibit hitchhiking-like behaviors in different organisms and contexts. In *S. cerevisiae*, mammalian neurons, and *U. maydis*, ribonucleoprotein complexes are tethered to and co-transported with different membrane-bound compartments (Göhre *et al*., 2012; Jansen *et al*., 2014; Haag *et al*., 2015; Cioni *et al*., 2019; Liao *et al*., 2019). In *U. maydis*, polysomes associate with the RNA-binding protein Rrm4, which interacts with the EE-associated protein Upa1 (Pohlmann *et al*., 2015). In neurons, ANXA11 links RNA granules to lysosomes (Liao *et al*., 2019). Membrane-bound organelles, including peroxisomes, also hitchhike on motile EEs in both *U. maydis* and *A. nidulans* (Guimaraes *et al*., 2015; Salogiannis *et al*., 2016). A genetic screen in *A. nidulans* identified the protein PxdA as a critical mediator of peroxisome hitchhiking (Salogiannis *et al*., 2016). However, how PxdA links peroxisomes to EEs and whether other proteins are involved remain unclear. In *U. maydis*, lipid droplets (LDs) and the endoplasmic reticulum, as well as peroxisomes, hitchhike on EEs (Guimaraes *et al*., 2015), although *U. maydis* lacks a PxdA homolog (Steinberg, 2016).

Here, we took a three-pronged approach to further determine the mechanism and role of hitchhiking in *A. nidulans*. First, we screened other organelles including mitochondria, autophagosomes, and lipid droplets and found that PxdA was not required for their proper distribution, suggesting that PxdA is a specific regulator of peroxisomes. Surprisingly, we found that the movement and distribution of lipid droplets were independent of both PxdA and HookA. Second, we returned to our initial screen (Tan *et al*., 2014) that identified PxdA and identified two new PxdA alleles, one of which is a point mutation (R2044P) that disrupts the association of PxdA with EEs. Third, we performed immunoprecipitation followed by mass spectrometry and identified the DenA/Den1 phosphatase DipA as a PxdA interacting protein. We showed that DipA is recruited to EEs in a PxdA-dependent manner and is required for peroxisome motility. Our data suggest that PxdA and the DipA phosphatase regulate peroxisome hitchhiking on EEs.

## Results

### Lipid droplets move independently of PxdA and early endosomes in Aspergillus

We sought to determine if other organelles require PxdA for their proper distribution along *A. nidulans* hyphae. To accomplish this, we fluorescently tagged the endogenous copies of Atg8, Tom20, and Erg6 to visualize autophagosomes, mitochondria, and lipid droplets, respectively. To quantify organelle distribution in wild-type and *pxdA*Δ cells, we generated line scans along hyphae and quantified their fluorescence intensity. All three organelles showed similar distribution profiles in wild-type and *pxdA*Δ hyphae (Figure 1A-B and Supplemental Figure S1), suggesting that these organelles do not use PxdA-mediated hitchhiking for transport. Therefore, this data suggests that rather than being a universal hitchhiking linker, PxdA may specifically regulate peroxisome movement in *A. nidulans*.

**FIGURE 1:**
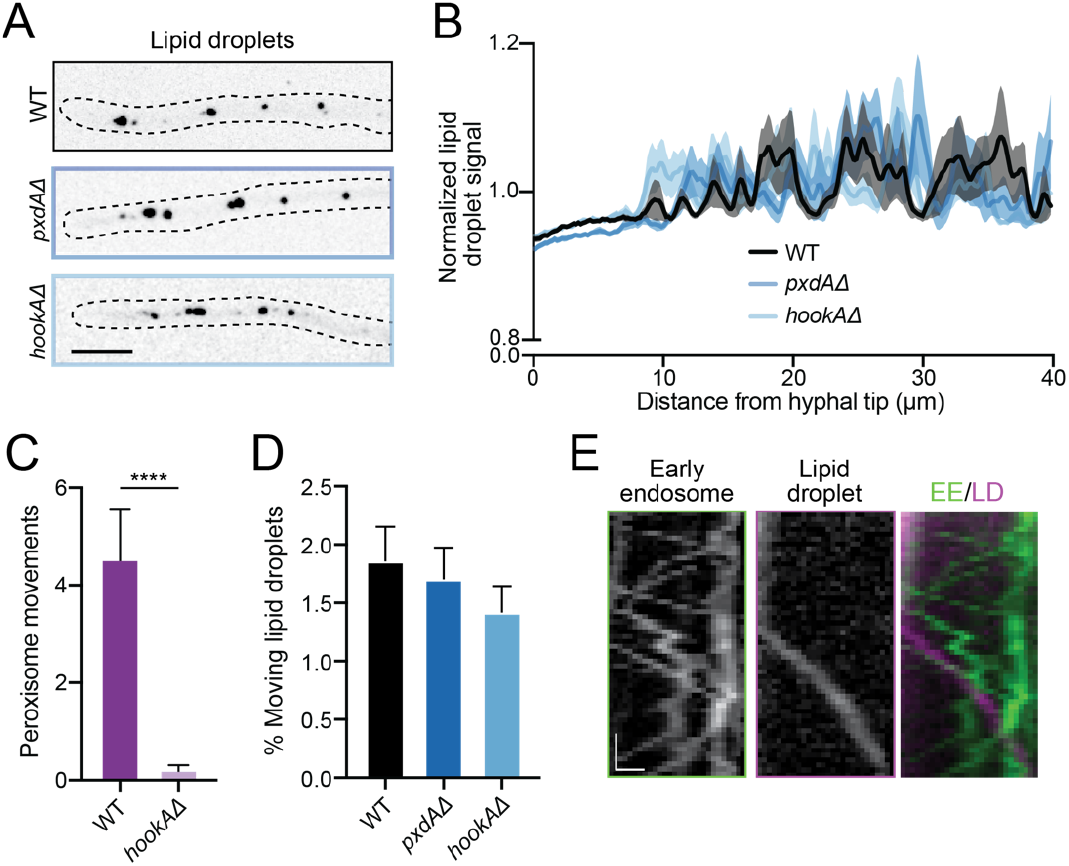
Lipid droplets do not hitchhike on early endosomes in *A. nidulans*. (A) Fluorescent micrographs of lipid droplets (Erg6/AN7146-mKate) along hyphae (outlined). Scale bar, 5 μm. (**B**) Lipid droplet distribution is quantified by fluorescence intensity line-scans. Normalized fluorescence intensity (solid lines) ± SEM (shading) is plotted as a function of distance from the hyphal tip. n = 55 (WT), 54 (*pxdAΔ*) and 45 (*hookAΔ*) hyphae. Genotype interaction is not significant by two-way ANOVA. **(C)** Bar graphs of peroxisome movements in WT and *hookAΔ* hyphae, quantified as the number of peroxisomes crossing a line drawn perpendicular and 10 μm away from the hyphal tip during a 30-sec time-lapse video. Data is mean ± SEM. n = 13 (WT) and 19 (*hookAΔ*) hyphae. ****p < 0.0001 (Mann-Whitney test). **(D)** Bar graphs displaying percent motility of lipid droplets (processive runs > 2.5 μm). Data is mean ± SEM. n = 14 (WT), 16 (*pxdAΔ*) and 32 (*hookAΔ*) field of views with > 1000 puncta analyzed per condition. Kruskal-wallis test was not-significant (n.s.). Also see Supplemental Movie S1. (**E)** Representative kymographs of EEs (GFP-RabA), lipid droplets and merged panel (EEs in green and lipid droplets (LD) in magenta). Scale bar, 1 μm. Time bar, 2 seconds. Also see Supplemental Movie S2.

Our findings that proper distribution of mitochondria and autophagosomes does not require PxdA is consistent with the lack of EE-mediated hitchhiking of mitochondria and late endosomes observed in *U*. maydis (Guimaraes *et al*., 2015). On the other hand, a PxdA-independent effect on the distribution of lipid droplets was unexpected given that they hitchhike on EEs in *U. maydis*. To further explore this, we abolished EE motility by deleting the EE-adaptor *hookA* (*hookA*Δ). While peroxisome motility was severely disrupted in *hookA*Δ cells, neither the distribution nor percentage of moving lipid droplets was affected in *hookA*Δ or *pxdA*Δ cells compared to wild-type cells (Figure 1A-D). Furthermore, though a small subset of lipid droplets (1.85 ± 1.02% [s.d.]) undergo long-distance, processive movement (runs greater than 2.5 μm; Supplemental Movie S1), these runs were not co-localized with EEs (Figure 1E, Supplemental Movie S2). Consistent with the distribution data, we find no differences in the run-length of lipid droplets and only slight differences in velocity amongst wild-type, *pxdA*Δ, or *hookA*Δ strains (Supplemental Figure S2). Taken together, this data suggests that lipid droplets do not hitchhike on EEs in *A. nidulans*.

### Identification of novel pxdA alleles

Since our data suggest that PxdA specifically regulates peroxisome motility, we next sought to further understand the mechanisms of PxdA-mediated hitchhiking of peroxisomes. To accomplish this, we returned to candidates from a previous mutagenesis screen established to identify novel regulators required for the microtubule-based transport of peroxisomes, EEs and/or or nuclei (Tan *et al*., 2014). In a follow-up study, we characterized two mutant strains with peroxisome-specific defects that mapped to two independent alleles in the gene *AN1156/pxdA*, which led to early stop codons (PxdA_S201stop_ and PxdA_Q846stop_) in the protein (Figure 2A) (Salogiannis *et al*., 2016). Here, we sequenced two additional mutants that showed defects in peroxisome movement and positioning. Both mutants were also mapped to *pxdA*. One of the new alleles created a stop codon at Q1201 (PxdA_Q1201stop_; Figure 2A), while the other allele, R2044P, was a single-residue mutation located in the coiled-coil 3 (CC3) domain of PxdA (PxdA_R2044P_; Figure 2A and Supplemental Figure S3A). Both new alleles showed defects in peroxisome motility but had normal EE motility and nuclear distribution (Figure 2B-E). We chose to focus on the *pxdA*_*R2044P*_ allele as the CC3 domain of PxdA is located within a region of the protein that is necessary and sufficient for PxdA to associate with early endosomes (Salogiannis *et al*., 2016).

**FIGURE 2:**
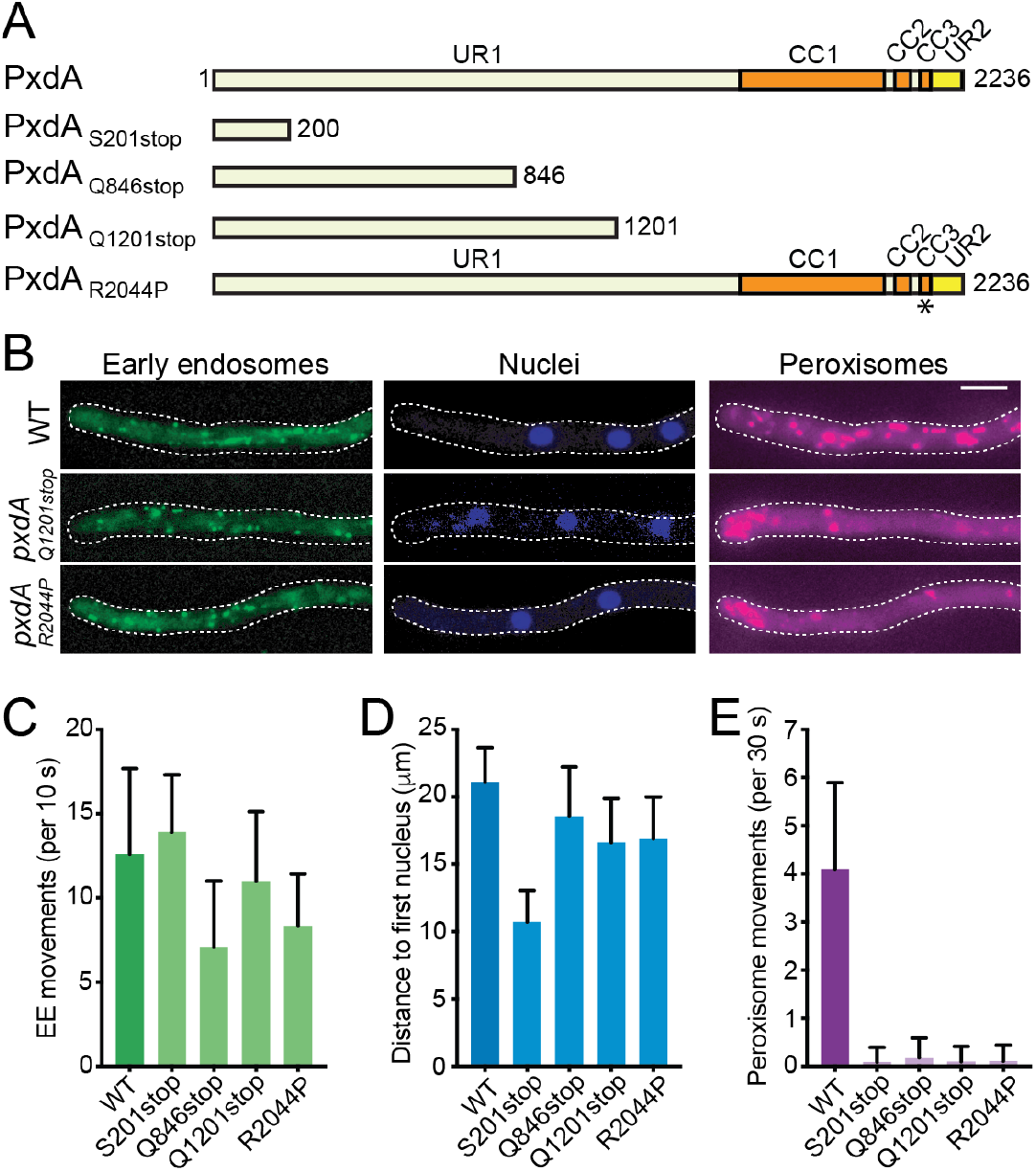
Novel *pxdA* alleles regulate peroxisome movement and distribution. **(A)** Schematic of PxdA domain organization and novel PxdA mutants. PxdA_S201stop_, PxdA_Q846stop_, and PxdA_Q1201stop_ mutations create stop codons that result in truncated proteins, while the PxdA_R2044P_ mutation is in the CC3 domain of PxdA (indicated by asterisk). **(B)** Representative images of EE (GFP-RabA), nuclear (HH1-BFP), and peroxisome (mCherry-PTS1) distribution in wild-type, PxdA_Q1201stop_ and PxdA_R2044P_ expressing hyphae. Scale bar, 5 μm. **(C-E)** Quantification of movement of EEs (C), distribution of nuclei (D), and movement of peroxisomes (E) in PxdA_Q1201stop_ and PxdA_R2044P_ compared to previously identified PxdA mutants (Tan *et al*., 2014; Salogiannis *et al*., 2016). EE and peroxisome movement is quantified as the number of EEs or peroxisomes crossing a line drawn perpendicular and 10 μm away from the hyphal tip during a 10- or 30-sec time-lapse video, respectively. Data is mean ± SD. n = 10 (WT), 11 (PxdA_Q846stop_),11 (PxdA_S201stop_), 10 (PxdA_Q1201stop_) and 9 (PxdA_R2044P_) hyphae. *p < 0.05, ****p< 0.0001 (Krustal-Wallis test with Dunn’s multiple comparisons test compared to WT strain).

### A point mutation in PxdA decreases its ability to associate with early endosomes

As the R2044P mutation is present within the CC3 domain, we hypothesized that this mutation would affect the ability of PxdA to associate with EEs. To test this, we created a fluorescently tagged version of PxdA_R2044P_ at the endogenous *pxdA* locus. Replacement of endogenous PxdA with PxdA_R2044P_-mTagGFP2 (PxdA_R2044P_-GFP) resulted in a significant decrease in peroxisome motility compared to a strain expressing wild-type PxdA-GFP (Figure 3A and Supplemental Movie S3). We then examined the localization and dynamics of PxdA_R2044P_-GFP compared to wild-type PxdA-GFP. PxdA_R2044P_-GFP showed a more diffuse localization, with few moving foci (Figure 3B, and Supplemental Movie S4), suggesting a defect in PxdA_R2044P_ association with EEs. Supporting this, we found that while many wild-type PxdA-GFP runs colocalized with EEs, there was little colocalization between PxdA_R2044P_-GFP and EEs (Figure 3C-D). A Western blot revealed that PxdA_R2044P_ was expressed at similar levels as the wild-type protein (Supplemental Figure S3B). Thus, a single mutation in the CC3 domain of PxdA is capable of disrupting PxdA’s association with EEs, resulting in decreased peroxisome hitchhiking.

**FIGURE 3:**
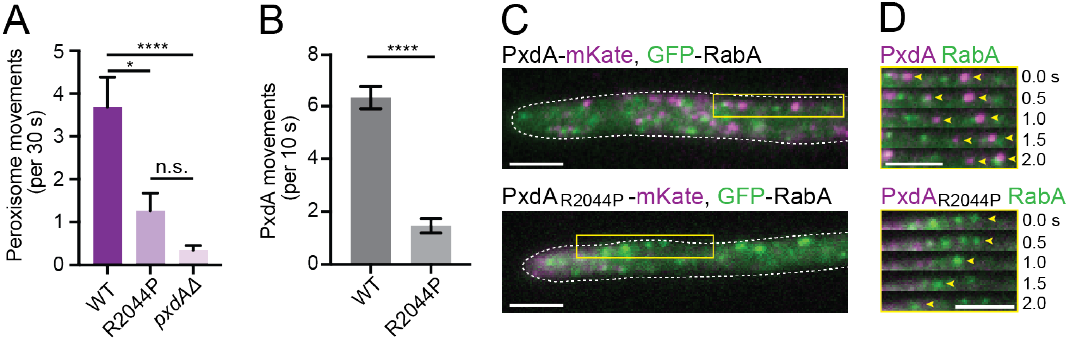
A point mutation in the CC3 domain of PxdA decreases PxdA association with early endosomes. **(A)** Bar graphs of peroxisome movements in WT, PxdA_R044P_, and *pxdAΔ* hyphae, quantified as the number of peroxisomes crossing a line drawn perpendicular and 10 μm away from the hyphal tip during a 30-sec time-lapse video. Data is mean ± SEM. n = 22 (WT), 27 (PxdA_R044P_), and 38 (*pxdAΔ*) hyphae. *p = 0.114, ****p < 0.0001 (Krustal-Wallis test with Dunn’s multiple comparisons test). Also see Supplemental Movie S3. **(B)** Bar graphs of PxdAWT-mTagGFP2 and PxdAR2044P-mTagGFP2 movements quantified as the number of PxdA puncta crossing a line drawn perpendicular and 10 μm away from the hyphal tip during a 10-sec time-lapse video. Data is mean ± SEM. n = 17 (WT), 17 (PxdA_R044P_) hyphae. ****p < 0.0001 (Mann Whitney test). Also see Supplemental Movie S4. **(C)** Representative images of hyphae expressing mTagGFP2-RabA and wild-type PxdA-mKate or mutant PxdA_R044P_-mKate. Scale bar, 5 μm. **(D)** Representative time-lapse stills from region indicated by yellow box in (C), demonstrating co-localization of EEs (GFP-RabA) with wild-type PxdA but not PxdA_R044P_. Yellow arrowheads denote processively-moving EEs (GFP-RabA). Scale bars, 5 μm.

### PxdA interacts with the DipA phosphatase on EEs

Our screen identified four alleles of *pxdA*, but no additional genes involved in hitchhiking. To identify other proteins that might be involved in hitchhiking, we immunoprecipitated PxdA followed by mass spectrometry. Specifically, we tagged PxdA at the endogenous locus with a hemagglutinin (HA) tag and performed immunoprecipitations with lysates from this strain or an untagged wild-type (WT) strain. PxdA resolves at approximately 250 kDa on SDS-PAGE gels when lysates are prepared under denaturing conditions (Salogiannis *et al*., 2016). However, when lysates are prepared under non-denaturing conditions, PxdA is partially degraded (Figure 4A and Supplemental Figure S3C). Despite this degradation, we could detect an additional faster migrating band (between 75-100 kDa) by Sypro stain (Figure 4A, right panel), which was present in the HA immunoprecipitations, but not the control. Mass spectrometry analysis revealed this band to be the ***D***enA/DEN1 ***i***nteracting ***p***hosphatase (DipA). DipA is a metallophosphatase that interacts with and regulates the stability of the deneddylase DenA/DEN1 on motile puncta in the cytoplasm, a process important for asexual fungal development (Schinke *et al*., 2016).

**FIGURE 4:**
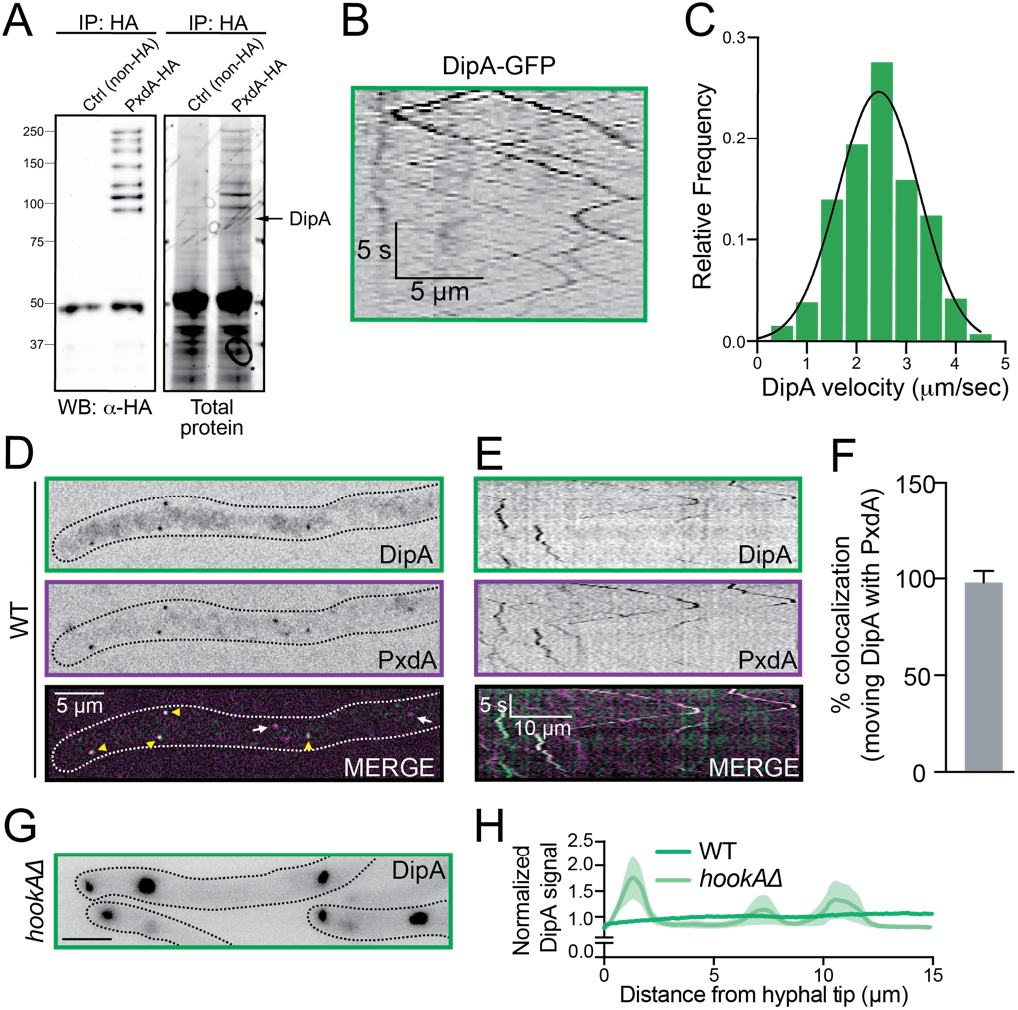
DipA associates with PxdA-marked endosomes. **(A)** Anti-HA western blot (left) and Sypro Ruby-stained SDS-PAGE gel (right) of lysates incubated with HA-conjugated agarose from WT or PxdA-HA expressing hyphae. Mass spectrometry identified DipA (indicated by arrow) from two biological replicates. **(B)** Representative kymograph of DipA-GFP movement. Also see Supplemental Movie S5. **(C)** Histogram of DipA-GFP velocities calculated from kymographs as in (B). Mean velocity is 2.44 ± 0.77 (SD) μm/sec. n = 257 moving events. **(D)** Colocalization of DipA-GFP and PxdA-mKate along a hypha. **(E)** Representative kymograph of DipA and PxdA co-movement. **(F)** Quantification of the percent overlap of DipA colocalized with PxdA. Mean % overlap is 97.88 ± 6.01 (SD). n=8 kymographs. **(G)** Representative image of DipA-GFP distribution in *hookAΔ* hyphae. **(H)** DipA-GFP distribution is quantified by fluorescence intensity line-scans of fluorescently-tagged organelles in wild-type or *hookAΔ* hyphae. Mean fluorescence intensity (solid lines) ± SEM (shading) is plotted as a function of distance from the hyphal tip. n=3 (WT) and 7 (*hookAΔ*) hyphae.

We next sought to determine where DipA is localized, how it interacts with PxdA, and whether it regulates peroxisome movement. To accomplish this, we constructed two DipA strains: (1) an endogenously tagged DipA with two tandem copies of TagGFP2 at its carboxy-terminus (DipA-GFP) to assess its localization, and (2) a DipA deletion (*dipA*Δ) strain to assess its cellular function. As previously reported, the *dipA*Δ strain exhibits perturbed colony growth (Supplemental Figure S4A) (Schinke *et al*., 2016). Colonies from the DipA-GFP strain grew normally (Supplemental Figure S4A), suggesting that the GFP tag does not adversely affect protein function. We performed live-cell imaging of DipA-GFP and found that DipA localizes to highly motile puncta (Figure 4B, and Supplemental Movie S5) that have a similar velocity to PxdA puncta and EEs (Figure 4C) (Abenza *et al*., 2009; Egan *et al*., 2012b; Salogiannis *et al*., 2016). Consistent with this, the vast majority of motile DipA puncta colocalize with PxdA (Figure 4D-F). Furthermore, in a *hookAΔ* strain in which EEs accumulate near the hyphal tip (Zhang *et al*., 2014), DipA displays similar accumulation (Figure 4G-H), suggesting that DipA associates with EEs along with PxdA.

Since PxdA and DipA interact and colocalize on the same motile EEs, we wondered if PxdA required DipA for its localization or vice versa. To test this, we first imaged the dynamics of PxdA in the absence of DipA (*dipA*Δ). We find that PxdA displays similar localization (Figure 5A) and motility in wild-type versus *dipA*Δ cells (Figure 5B and C and Supplemental Movie S6). In contrast, we find that DipA is largely cytosolic in *pxdA*Δ hyphae (Figure 5D) and exhibits drastically reduced long-range movement (Figure 5E-F and Supplemental Movies S7). There are no differences in DipA protein expression levels in *pxdA*Δ vs WT strains (Supplemental Figure S4C). These results suggest that PxdA is required for the recruitment of DipA to EEs.

**FIGURE 5:**
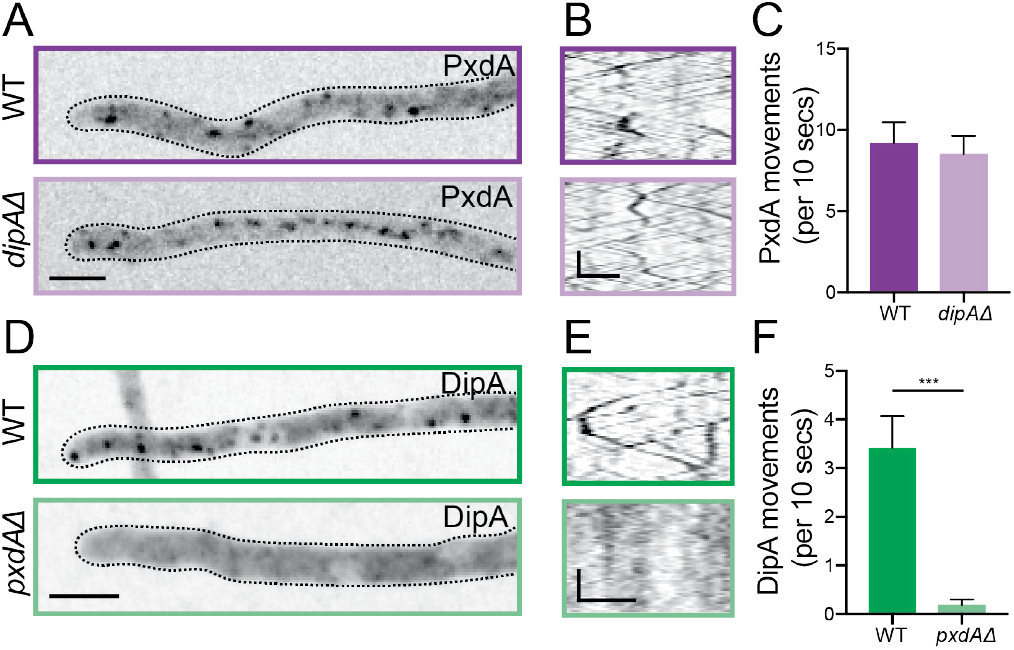
PxdA is required for DipA localization on moving foci. **(A)** Representative micrograph of GFP-PxdA puncta along a WT or *dipAΔ* hypha. Scale bar, 5 μm. **(B)** Representative kymograph of PxdA movement in a WT (top) or *dipAΔ* (bottom) background. Scale bars, 2.5 μm (horizontal), 5 s (vertical). Also see Supplemental Movie S6. **(C)** Bar graph of the flux of PxdA movements in WT and *dipAΔ* hyphae calculated as the number of PxdA foci crossing a line drawn perpendicular and 10 μm away. n=6 (WT) and 10 (*dipAΔ*) hyphae. **(D)** Representative micrograph of DipA-GFP puncta along a WT or *pxdAΔ* hypha. Scale bar, 5 μm. **(E)** Representative kymograph of DipA movement in a WT (top) or *pxdAΔ* (bottom) background. Scale bars, 2.5 μm (horizontal), 5 s (vertical). Also see Supplemental Movie S7. **(F)** Bar graph of the flux of DipA movements in WT and *pxdAΔ* hyphae calculated as the number of DipA foci crossing a line drawn perpendicular and 10 μm away. n=25 (WT) and 11 (*pxdAΔ*) hyphae. ***p=0.0002 (Mann Whitney test).

### DipA regulates peroxisome movement

Based on our observations thus far, we hypothesized that DipA is required for peroxisomes to hitchhike on moving EEs. To test this, we examined peroxisome and EE distribution in a *dipA*Δ strain. Peroxisomes in *dipAΔ* hyphae displayed a slight, but significant, accumulation near the hyphal tip (Figure 6A-B), while EEs were distributed normally (Figure 6C-D). The *dipA*Δ strain also showed a drastic reduction in peroxisome movement, but normal movement of EEs (Figure 6E-F, and Supplemental Movies S8 and S9). Therefore, the *dipA*Δ strain phenocopies the *pxdA*Δ strain, consistent with our observation that both proteins associate with the same subset of EEs.

**FIGURE 6:**
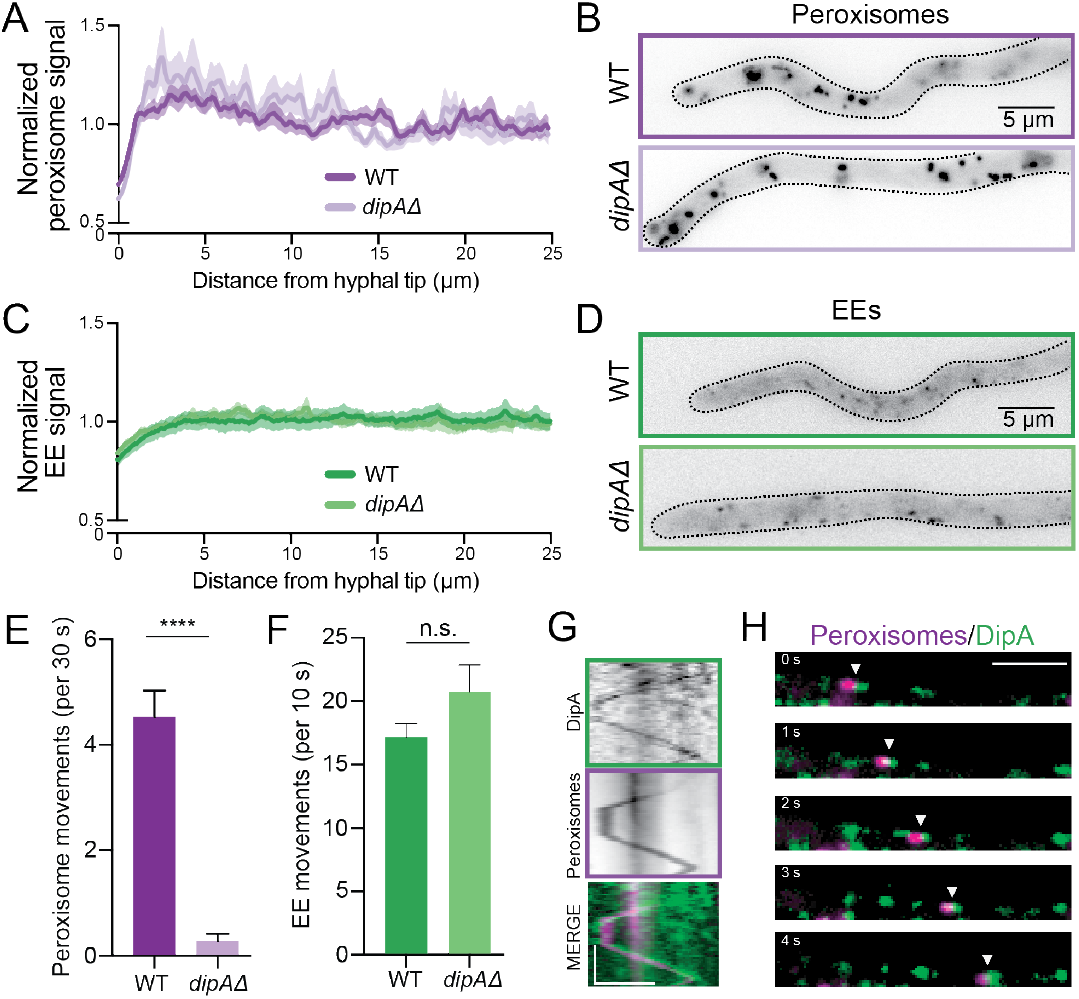
DipA regulates the movement and distribution of peroxisomes. **(A)** Distribution of peroxisomes along WT and *dipAΔ* hyphae. Data is normalized mean intensity ± SEM. n=22 (WT) and 20 (*dipAΔ*) hyphae. Distribution comparing genotypes is significant (p<0.0001) by two-way ANOVA. **(B)** Representative micrographs of peroxisomes along WT and *dipAΔ* hyphae. **(C)** Distribution of EEs along WT and *dipAΔ* hyphae. n=7 (WT) and 6 (*dipAΔ*) hyphae. Distribution comparing genotypes is not significant by two-way ANOVA. **(D)** Representative micrographs of EEs along WT and *dipAΔ* hyphae. (**E)** Bar graph of the flux of peroxisome movements in WT and *dipAΔ* hyphae calculated as the number of peroxisomes crossing a line drawn perpendicular and 10 μm away (*dipAΔ)* hyphae). ****p < 0.0001, Mann-Whitney test. Also see Supplemental Movie S8. **(F)** Bar graph of flux of EE movements in WT and *dipAΔ* hyphae calculated as in Figure 2C. n= 7 (WT) and 16 (*dipAΔ*) hyphae. Also see Supplemental Movie S9. **(G-H)** Kymographs (G) and time-lapse stills (H) of DipA-GFP with moving peroxisomes. Scale bars, 5 μm (horizontal) and 5 s (vertical). Also see Supplemental Movie S10.

Finally, we examined the localization of DipA in reference to hitchhiking peroxisomes. During peroxisome hitchhiking, PxdA-labeled EEs colocalize at the leading edge of moving peroxisomes (Guimaraes *et al*., 2015; Salogiannis *et al*., 2016), suggesting that PxdA and EEs might dictate the directionality of moving peroxisomes. We were curious if moving DipA foci interacted with peroxisomes in a similar manner. We first monitored the flux of peroxisomes in the DipA-GFP strain and found that tagging DipA does not perturb peroxisome movement (Supplemental Figure S4B). Next, using simultaneous two-color imaging of DipA and peroxisomes, we found that motile DipA localizes to the leading edge of peroxisomes (Figure 6G-H), similar to our previously published data on PxdA (Salogiannis *et al*., 2016). In further support of this, DipA shifts to the leading edge of peroxisomes concurrent with rapid directional switches in peroxisome movement (Figure 6G and Supplemental Movie S10). In summary, we find that DipA localizes to PxdA-labeled EEs and is critical for peroxisome hitchhiking.

## Discussion

Proper cargo transport is critical for many cell types, including filamentous fungi, in which cellular cargos are evenly distributed along their hyphae. Hitchhiking is a mode of transport whereby one type of cargo achieves motility via attachment to another type of motile cargo. We previously found that the protein PxdA is required for peroxisomes to hitchhike on EEs in *A. nidulans*. In this study, we find that PxdA specifically regulates peroxisomes, whereas lipid droplet, mitochondria, and autophagosome distribution are not affected in the absence of PxdA. We identified two novel mutant alleles of PxdA, including a single residue mutation (PxdA_R2044P_) that disrupts PxdA association with EEs. We also identified a novel regulator of peroxisome hitchhiking, the phosphatase DipA. DipA is required for peroxisome motility, colocalizes with PxdA, and its association with EEs requires PxdA. Together, these data provide further insight into factors that affect peroxisome hitchhiking in *A. nidulans*.

### Lipid droplets do not hitchhike on early endosomes in Aspergillus

Unlike in *U. maydis*, LDs do not hitchhike on EEs in *A. nidulans*. We conclude this for the following reasons: First, the movement and distribution of LDs is similar to wild-type in both *pxdA*Δ and *hookA*Δ strains, the latter of which completely abolishes EE motility. Second, we did not detect any co-transport between EEs and LDs. Third, the velocity of LDs in *A. nidulans* is less than 1 μm/sec, which is much slower than the velocity of EEs, PxdA, and DipA foci (~2.5 μm/sec). In fact, in *U. maydis*, the velocity of LDs was reported to be much faster (~1.8 μm/sec) and very similar to the velocity of peroxisomes (Guimaraes *et al*., 2015). These findings suggest that that *U. maydis* and *A. nidulans* potentially have very different modes of organelle distribution and transport. Indeed, the *A. nidulans* peroxisome hitchhiking tether PxdA is not conserved in *U. maydis*, and the regions of the *U. maydis* ribonucleoprotein tether Rrm4/Upa1 required for hitchhiking are not conserved in *A. nidulans* (Pohlmann *et al*., 2015), suggesting that distinct hitchhiking machineries have evolved in *U. maydis* and *A. nidulans*. Future work will be needed to identify how the transport and distribution of lipid droplets and other cargos are regulated in different filamentous fungi.

### A single residue mutation in PxdA disrupts its association with early endosomes

We previously showed that the CC2 and CC3 domains of PxdA are required for its association with EEs (Salogiannis *et al*., 2016). Here, we found that a PxdA mutant allele (*pxdA*_*R2044P*_) disrupts PxdA association with EEs via a single amino acid change in coiled-coil region CC3. This mutation may disrupt PxdA’s ability to interact with a binding partner on the EE surface or with the EE membrane directly. The PxdA_R2044P_ mutant will be an important tool in the future to characterize how PxdA associates with EEs and whether PxdA directly or indirectly links EEs to hitchhiking cargos.

### DipA is a regulator of peroxisome hitchhiking

Our mass spectrometry experiment identified the phosphatase DipA as a PxdA interactor. DipA associates with the PxdA-bound population of EEs and is required for peroxisomes to hitchhike on EEs. Additionally, DipA requires PxdA to associate with EEs. DipA is a phosphatase whose function was previously linked to regulating the phosphorylation status and degradation of DenA/Den1, a protein involved in asexual spore formation (Christmann *et al*., 2013; Schinke *et al*., 2016). DenA is co-transported with DipA (Schinke *et al*., 2016). Given the similar co-transport of both DenA and PxdA with DipA, it is possible that DipA also regulates PxdA by modulating PxdA’s phosphorylation state. Future work will determine whether PxdA is phosphorylated, whether its phosphorylation state is regulated by DipA, and whether DenA is required for hitchhiking.

Though we do not understand how the PxdA-DipA complex links to peroxisomes, it is also possible that DipA regulates the phosphorylation state of other proteins in the hitchhiking machinery, such as peroxisome-associated proteins. Alternatively, DipA may not regulate the phosphorylation state of the hitchhiking machinery at all but may instead link PxdA to peroxisomes. Whether DipA links PxdA to peroxisomes or whether DipA regulates hitchhiking in other ways remains to be determined.

## Materials and Methods

### Fungal growth conditions

*A. nidulans* strains were grown in yeast extract and glucose (YG) medium or 1% glucose minimal medium(Nayak *et al*., 2006), supplemented with 1 mg/ ml uracil, 2.4 mg/ ml uridine, 2.5 μg/ ml riboflavin, 1 μg/ml para-aminobenzoic acid, and 0.5 μg/ ml pyridoxine when required. Glufosinate (Sigma) for *bar* selection was used at a final concentration of 700 μg/ mL (Straubinger *et al*., 1992).

For imaging of germlings, spores were resuspended in 0.5 mL 0.01% Tween-80 solution. The spore suspension was diluted at 1:1,000 in liquid minimal medium containing appropriate auxotrophic supplements. The spore and media mix (400 μL) was added to an eight-chambered Nunc Lab-Tek II coverglass (ThermoFisher) and incubated at 30°C for 16-20 hours before imaging. For imaging of mature hyphae, spores were inoculated on minimal medium plates containing the appropriate auxotrophic supplements and incubated at 37°C for 12-16 hours. Colonies were excised from agar plates and inverted on Lab-Tek plates for imaging. For biochemistry, spores were inoculated in YG medium containing the appropriate auxotrophic supplements for 16-20 hours at 37°C shaking at 200 rpm.

### Plasmid and strain construction

Strains of *A. nidulans* used in this study are listed in Table S1. All strains were confirmed by a combination of PCR and/or sequencing from genomic DNA isolated as previously described(Lee and Taylor, 1990) and in some cases by Western blot analysis. For all plasmids, PCR fragments were inserted into the Blue Heron Biotechnology pUC vector at 5’ EcoRI and 3’ HindIII restriction sites using Gibson isothermal assembly(Gibson *et al*., 2009). All plasmids were confirmed by Sanger sequencing. Initial strains were created by homologous recombination with linearized DNA to replace the endogenous gene in strains lacking *ku70* with *Afribo (Aspergillus fumigatus Ribo), AfpyrG* (*Aspergillus fumigatus pyrG*), or *Afpyro* (*Aspergillus fumigatus pyro*), or *bar* as selectable markers (Straubinger *et al*., 1992; Nayak *et al*., 2006). Additional strains were created using genetic crossing as previously described (Todd *et al*., 2007). The strategies for transforming *hookA*Δ, *pxdA*Δ, PxdA-HA, PxdA-mTagGFP2, and PxdA-mKate2 strains, as well as the strains containing fluorescently labeled EEs, peroxisomes, and nuclei, have been previously described (Tan *et al*., 2014; Salogiannis *et al*., 2016). The strategies for DNA constructs and strains created for the current study are as follows:

#### PxdA_R2044P_-GFP/mKate

To construct this plasmid, carboxy-terminal codon-optimized fluorescent protein tags, either mTagGFP2 (Subach *et al*., 2008) or mKate2 (Shcherbo *et al*., 2009) were preceded by a GA(x4) linker and followed by *AN1156/pxdA*’s native 3’ UTR with flanking 1kb upstream and downstream homologous recombination arms. To introduce the Q2044P mutation, the upstream 1kb recombination arm was amplified by PCR from genomic DNA (gDNA) isolated from RPA568 (mutagenized strain harboring the Q2044P mutation) using the oligos 5’-GTCACGACGTTGTAAAACGACGGCCAGTGCCTTCAGGAGCGAGTGGCGCATCCGAAG-3’ and 5’-CTGACATTGCACCTGCACCTGCACCTGCACCTGCTATTTGTGGGCCAAAGGGACCGTCGG-3’. To amplify the linearized DNA used for targeting via transformation, the oligos 5’-CCTTCAGGAGCGAGTGGCGCATCTC-3’ and 5’-GAC GTTGACACTTCGTGCTAGAACT-3’ were used.

#### GFP-PxdA

This N-terminal GFP-tagged PxdA construct was used for data collected in Figure 5, A-C. For ease of cloning, this construct lacks the first 816 aa of PxdA (PxdA(Δ1-816) starts with 5’-CAGCAGGTCCCGGTTCCAC-3’ for reference), but is functional since it has no effect on peroxisome motility and distribution (data not shown) (Salogiannis *et al*., 2016). Codon-optimized mTagGFP2 with a GA(x4) linker was followed by the PxdA(Δ1-816) fragment (4260 bp), its native 3’ UTR and an *AfpyrG* cassette, and flanked by 1kb homologous recombination arms. To amplify the linearized DNA used for transformations, the oligos 5’-AGTCGACACGGAAGGTTGGTCAATC-3’ and 5’-GACGTTGACACTTCGTGCTAGAACT-3’ were used. To identify positive clones with better efficiency, we targeted the linear PxdA(Δ1-816)-*AfpyrG*-containing DNA into a *pxdA*Δ strain (RPA921) where the endogenous pxdA locus was replaced with *Afribo*. Using this strategy we positively selected clones that (1) grow on minimal media plates lacking supplemented uradine-uracil and (2) are unable to grow on plates lacking riboflavin. From there, we sequenced verified this strain and used genetic crosses to generate additional strains.

#### dipAΔ

This strain was originally previously published elsewhere (Schinke *et al*., 2016), but it was remade for this study using a different strategy. Linearized DNA used for targeting was amplified with 5’-ATGAGACGAACCTGGCCATCAAGGC-3’ and 5’-TTATGCCATGTTGCAGGTGGAA CA-3’ by fusion PCR (Szewczyk *et al*., 2006) with fragments for a selectable marker flanked by upstream and downstream homologous recombination arms around the *AN10946/dipA* genomic locus.

#### DipA-GFP

A tandem 2x-mTagGFP2 preceded by a GA(x4) linker and followed by *AN10946/dipA*’s native 3’ UTR with flanking 1kb upstream and downstream homologous recombination arms. This plasmid was digested with the Pci restriction enzyme (cut sites in the 5’ and 3’ homologous arms) to linearize DNA for targeting.

#### AN7146-mKate (lipid droplets)

To visualize endogenously-labeled lipid droplets, we fluorescently tagged AN7146 (a putative S-adenosyl-methionine delta-24-sterol-C-methyltransferase). AN7146 is a homolog of Erg6 in Ustilago maydis and used as a lipid droplet marker in a previous study(Guimaraes et al., 2015). To create this plasmid, C-terminal codon-optimized mKate2 was preceded by a GA(x4) linker and followed by *AN7146*’s native 3’ UTR with flanking 1kb upstream and downstream homologous recombination arms. The oligos 5’-GTTGTTCAAAACCCTTGGGAAATTTG-3’ and 5’-TCCAGTTGAAGTACACTACACATTCG-3’ were used to amplify the linearized DNA fragment used for targeting.

#### GFP-AtgH (autophagosomes)

To visualize autophagosomes, we fluorescently tagged the functional homolog of Atg8, *atgH/AN5131*. To construct this plasmid, the codon-optimized mTagGFP2 followed by a GA5 linker and the entire *atgH* locus (including its native 3’UTR) was flanked by homologous recombination arms. The oligos 5’-CTGTAAATTCTTTCTTGCCC-3’ and 5’-CTTTGCCGTCGTATCGACC-3’ were used to PCR amplify the targeting construct used for transformation.

#### Tom20-GFP (mitochondria)

To visualize mitochondria, we fluorescently tagged the mitochondrial outer membrane component Tom20 (Suresh *et al*., 2017), *AN0559*. To construct this plasmid, a codon-optimized 2xmTagGFP2 was preceded by a GSGSG linker and followed by *AN0559*’s native 3’ UTR with flanking 1kb upstream and downstream homologous recombination arms. The oligos 5’-GGGCGGCGATTTTTGCTGCGAAGAG-3’ and 5’-CGGATTTGCCGTCAACGGCGTTGGT-3’ were used to amplify the linearized DNA fragment used for targeting.

### Whole-genome sequencing

The general workflow for whole genome sequencing has been previously described (Tan *et al*., 2014). The mutagenized strains, RPA568 and RPA639, were backcrossed to the parental RPA520 strain while tracking the presence of the peroxisome accumulation and motility phenotypes in order to (1) reduce the number of background mutations and (2) to ensure the phenotype likely arose from a single gene (similar to Tan *et al*. 2014). Genomic DNA (gDNA) was prepared from a pool of at least five individual backcrossed strains and treated with RNase prior to a final cleanup by phenol/chloroform extraction. gDNA was sheared with the Covaris S2 sonicator, and samples were run on a D1000 Tapestation to evaluate shearing success. Samples were normalized to the same input and prepped using KAPA HTP Library Prep Reagents using an Apollo 324 robot. This involved End-Repair, A-Tailing, and Adapter Ligation, followed by amplification and barcoding (similar to [9]). A bead-based cleanup removed primer-dimers and adapter-dimers. The final library products were run on a High Sensitivity D1000 Screentape and qPCR was performed using the KAPA Library Quantification kit. The libraries were then pooled and loaded onto an individual lane of the HiSeq 2500 Rapid flowcell, with a read length of ~50 base pairs. Data sets were sorted by barcode using Python software as previously described and the data were aligned using the *A. nidulans* reference genome FGSC_A4. Genomic data was sorted for genomic feature (i.e. exon, intron) and mined specifically at the *AN1156/pxdA* locus for high quality (> 5 times in the mutant data set and zero in the backcrossed WT mutagenized strain) and uncommon (i.e., not in the WT backcrossed strain) nonsynonymous mutations.

### Imaging acquisition

All images were collected at room temperature. Imaging experiments in Figure 2, Figure 3, Figure 4B,C,G,H and Figure 6A-F, Supplementary Figure 1, and Movies S2-4, S6, S8-10 were acquired using an inverted microscope (Nikon, Ti-E Eclipse) equipped with a 60x and 100x 1.49 N.A. oil immersion objective (Nikon, Plano Apo), and a MLC400B laser launch (Agilent), with 405 nm, 488 nm, 561 nm and 640 nm laser lines. Excitation and emission paths were filtered using single bandpass filter cubes (Chroma). For two-color colocalization imaging in Figure 2, the emission signals were further filtered and split using W-view Gemini image splitting optics (Hamamatsu). Emitted signals were detected with an electron multiplying CCD camera (Andor Technology, iXon Ultra 888). Illumination and image acquisition were controlled with NIS Elements Advanced Research software (Nikon), and the xy position of the stage was controlled with a ProScan linear motor stage controller (Prior). Simultaneous two-color time-lapse images in Figure 4D-F and Figure 6G-H were collected using a Plan Apo TIRF 60x/1.49 oil immersion objective on the Deltavision OMX Blaze V4 system (GE Healthcare). GFP and mKate2/mCherry were excited simultaneously with 488nm and 568nm diode laser lines, respectively. A BGR polychroic mirror was used to split emission light from fluorophores to different PCO Edge sCMOS cameras. Emission filters in front of both cameras was used to select appropriate wavelengths (528/48 and 609/37 for GFP and mKate/mCherry, respectively). Images were aligned with OMX image registration using softWoRx software and some images were deconvolved for display using the enhanced ratio method.

For imaging experiments in Figure 1A-D, Supplementary Figure 2, Figure 5, and Movies S1, S5, and S7, spinning disk confocal microscopy was performed using a Yokogawa W1 confocal scanhead mounted to a Nikon Ti2 microscope with an Apo TIRF 100x 1.49 NA objective. The scope was controlled via NIS Elements using the 488nm and 561nm lines of a six-line (405nm, 445nm, 488nm, 515nm, 561nm, and 640nm) LUN-F-XL laser engine and a Prime95B camera (Photometrics). Image channels were acquired using bandpass filters for each channel (525/50 and 595/50). Z-stacks were acquired using a piezo Z stage (Mad City Labs).

### Image and Data analysis

Line-scan distribution and flux measurements were calculated as previously reported (Salogiannis *et al*. 2016). For line-scan measurements, maximum-intensity projections of fluorescence micrographs and brightfield images (for hyphae) were obtained using ImageJ/FIJI (National Institutes of Health, Bethesda, MD). Brightfield images were traced starting from the hyphal tip, using the segmented line tool (line width 25), and traces were superimposed on the fluorescence micrographs to project the average fluorescence intensity. For normalization of line-scans, each condition’s average immunofluorescence intensity values at each point along the hyphae were normalized against that condition’s total baseline average. For flux measurements, the number of puncta crossing a line approximately 10 μm perpendicular to and from the hyphal tip was manually counted. All peroxisome flux data were calculated from 30-sec time-lapse videos, while all EE, DipA, and PxdA flux data were calculated from 10-sec time-lapse videos. DipA velocity (Figure 4C) and lipid droplet velocity (Supplementary Figure 2B) was analyzed using ImageJ. Specifically, maximum-intensity projections were generated from time-lapse sequences to define the trajectory of particles of interest. The segmented line tool was used to trace the trajectories and map them onto the original video sequence, which was subsequently resliced to generate a kymograph. The instantaneous velocities and run lengths of individual particles were calculated from the inverse of the slopes of kymograph traces. To calculate the percent motility of LDs (Figure 1), 3-min time-lapse movies of a field of view of mature hyphae (containing segments of ~5-15 cells) expressing fluorescently-labeled lipid droplets (AN7146-mKate) were imaged at 500 millisecond intervals. After background subtraction in ImageJ, moving LDs were initially assessed visually and kymographs were subsequently generated (as described above) from these moving puncta. An LD puncta was scored as moving if it exhibited a directed run (no pauses) greater than 2.5 μm during its movement. This number was divided by the total number of LD puncta in the field of view, which was calculated by using the ‘find maxima’ tool in ImageJ on the first frame of each movie. At least 50 LDs were analyzed per field of view and more than 1000 total LDs were analyzed per condition across three separate days. For distance to first nucleus measurements (Figure 2D), nuclei were thresholded in ImageJ and the segmented line tool was used to measure the distance from the hyphal tip to the edge of the first nucleus.

For DipA/PxdA colocalization measurements, DipA and PxdA kymographs were generated from simultaneous two-color time-lapse videos. DipA kymographs were thresholded in ImageJ, manually traced using the segmented line tools, and traces were then superimposed onto the PxdA kymograph to manually determine overlap. The fractional overlap (colocalization) of DipA with PxdA for each kymograph was calculated. Due to the weak and unreliable fluorescence signal of DipA in our time-lapse videos, we were unable to determine the fraction of PxdA that colocalized with DipA.

All statistical analyses were performed using Prism8 (GraphPad).

### Cell Lysis and Immunoprecipitations

Overnight cultures for biochemistry were strained in Miracloth (Millipore), flash frozen in liquid nitrogen, and ground with a mortar/pestle in the presence of liquid nitrogen. For Western blot analysis of DipA-GFP in WT and *pxdAΔ* strains (Supplemental Figure S4C), PxdA-GFP and PxdA_R2044P_-GFP comparisons (Supplemental Figure S3B) and Western blot analysis of PxdA-TurboID-3xFLAG strain in urea buffer (Supplemental Figure S3C), 250 uL of packed ground mycelia were resuspended in 500 uL boiling denaturing buffer (125 mM Tris-HCl pH 7.0, 8M Urea, 1mM EDTA pH7.0, 2% SDS, 10mM DTT, 10% beta-ME, 4% glycerol), rotated for 10 mins at room temperature (RT), spun at 20,000 x g for 10 mins at RT, and followed by addition of 1x Laemmli buffer (BioRad) to the supernatant. For western blot analysis of PxdA-TurboID-3xFLAG strain in RIPA buffer (Supplemental Figure S3C), 250 uL of packed ground mycelia were resuspended in 500 uL of RIPA buffer (50 mM Tris-HCl, pH 8.0; 150 mM NaCl, 1% (v/v) NP-40, 0.5% (w/v) sodium deoxycholate, 0.1% (w/v) SDS, 1 mM DTT) with added protease inhibitors (cOmplete Protease Inhibitor Cocktail, Roche; Switzerland) at 4°C rotated for 10 mins at room temperature (RT), spun at 20,000 x g for 10 mins at RT, and followed by addition of 1x Laemmli buffer (BioRad) to the supernatant. For anti-HA immunoprecipitations from WT and PxdA-HA strains used for mass spectrometry analysis (Figure 4A), all steps were performed at 4°C unless otherwise stated. Approximately 4 mLs packed ground mycelia were lysed in 10 mLs of modified RIPA buffer (150 mM NaCl, 50 mM Tris pH 8.0, 1% Triton X-100, 0.1% SDS) supplemented with protease inhibitor cocktail (Roche). Lysates were rotated for 30 mins, and centrifuged for 5 mins at 1000 x g. Supernatants were subjected to two additional 20,000 x g spins for 20 mins each. Agarose beads were added to clarified supernatants and rotated over-end for 1 hr to pre-clear any non-specific interactions. Pre-cleared lysate was incubated with HA-conjugated agarose beads (ThermoFisher) and rotated over-end for 1.5 hrs. Beads were washed 3x with modified RIPA buffer and were subsequently boiled in Laemmli buffer for 5 mins prior to running on an SDS-PAGE gel.

### Western blotting and mass spectrometry

All protein samples were resolved on 4-12% gradient SDS-PAGE gels (Life Technologies) for 60 mins. For Western blotting, gels were transferred to nitrocellulose for 3 hrs at 250 mA in 4°C. Blots were blocked with 5% milk in TBS-0.1% Tween-20 (TBS-T). Antibodies were diluted in 5% milk in TBS-T. Primary antibodies were incubated overnight at 4°C and horseradish peroxidase (HRP)-conjugated secondary antibodies (ThermoFisher) were incubated at RT for 1 hr. Rabbit anti-Tag(CGY)FP (Evrogen) was used at 1:1000 to detect mTagGFP2. Mouse anti-HA (Sigma) was used at 1:1000. Secondary antibodies were used at 1:10,000. Blots were rinsed 3x TBS-T for 10 mins after primary and secondary antibodies. Electrochemiluminescence was produced with HRP and Prosignal Pico (Prometheus) detection reagent and imaged on a ChemiDoc (BioRad) using Image Lab (v5.2.1.) software.

For Sypro protein staining and subsequent mass spectrometry of anti-HA immunoprecipitations (Figure 4A), SDS-PAGE gel was incubated in Sypro Red protein gel stain (ThermoFisher) at 1:5000 in 7.5% acetic acid for 45 mins. Band of interest was excised and submitted to the Taplin Mass Spectrometry Facility at Harvard Medical School. A before and after picture using Sypro Ruby detection on the ChemiDoc (BioRad) was obtained to ensure the correct bands were excised. Excised gel bands were cut into approximately 1 mm3 pieces. Gel pieces were then subjected to a modified in-gel trypsin digestion procedure (Shevchenko *et al*., 1996). Gel pieces were washed and dehydrated with acetonitrile for 10 min followed by removal of acetonitrile. Pieces were then completely dried in a speed-vac. Rehydration of the gel pieces was with 50 mM ammonium bicarbonate solution containing 12.5 ng/μl modified sequencing-grade trypsin (Promega, Madison, WI) at 4°C. After 45 min., the excess trypsin solution was removed and replaced with 50 mM ammonium bicarbonate solution to just cover the gel pieces. Samples were then placed in a 37°C room overnight. Peptides were later extracted by removing the ammonium bicarbonate solution, followed by one wash with a solution containing 50% acetonitrile and 1% formic acid. The extracts were then dried in a speed-vac (~1 hr) and stored at 4°C until analysis. On the day of analysis the samples were reconstituted in 5 - 10 μl of HPLC solvent A (2.5% acetonitrile, 0.1% formic acid). A nano-scale reverse-phase HPLC capillary column was created by packing 2.6 μm C18 spherical silica beads into a fused silica capillary (100 μm inner diameter x ~30 cm length) with a flame-drawn tip. After equilibrating the column each sample was loaded via a Famos auto sampler (LC Packings, San Francisco CA) onto the column. A gradient was formed and peptides were eluted with increasing concentrations of solvent B (97.5% acetonitrile, 0.1% formic acid). As peptides eluted they were subjected to electrospray ionization and then entered into an LTQ Orbitrap Velos Pro ion-trap mass spectrometer (Thermo Fisher Scientific, Waltham, MA). Peptides were detected, isolated, and fragmented to produce a tandem mass spectrum of specific fragment ions for each peptide. Peptide sequences (and hence protein identity) were determined by matching to the *A. nidulans* FGSC A4 protein database in Uniprot with the acquired fragmentation pattern by the software program, Sequest (Thermo Fisher Scientific, Waltham, MA). All databases include a reversed version of all the sequences and the data was filtered to between a one and two percent peptide false discovery rate.

## Author Contributions and Notes

JS, JRC, and SLR-P devised the experiments. JS and JRC performed all experiments with help from AA-M. NS shared unpublished findings about DipA. SLR-P supervised the research. JS, JRC, and SLR-P wrote and edited the manuscript, with feedback from NS.

## Acknowledgments

We thank Stephen A. Osmani for personal communication and sharing unpublished data on DipA and Thomas L. Schwarz for his guidance and supervision of J.S. for part of this project. We also thank the Nikon Imaging Centers at Harvard Medical School and UC San Diego for technical support and advice, Ross Tomaino and the Taplin Mass Spectrometry Facility at Harvard Medical School for technical support on mass spectrometry experiments, and the Bioinformatics Department at the Harvard T.H. Chan School of Public Health for assisting with whole genome sequencing data analysis. JS was funded for part of this work by the Charles King Trust Fellowship supported by the Charles H. Hood Foundation. JRC is funded by a postdoctoral fellowship from the National Institutes of Health (F32GM126692). SLR-P is an investigator of the Howard Hughes Medical institute and is also supported by R01GM121772.

