## Supplemental Information for "PxdA interacts with the DipA phosphatase to regulate endosomal hitchhiking of peroxisomes"

### SUPPLEMENTAL FIGURES

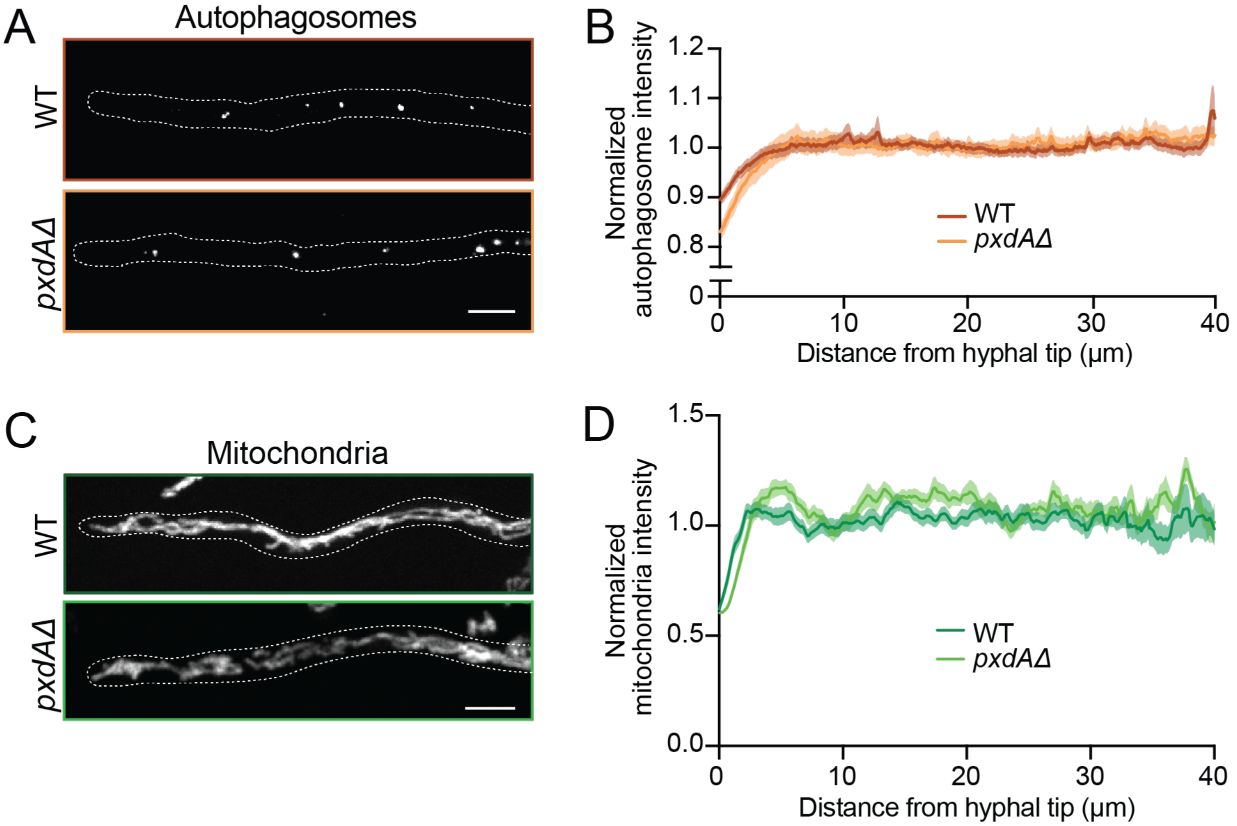


**Supplemental Figure S1: PxdA does not regulate the distribution of autophagosomes or mitochondria. (A)** Representative micrographs of wild-type or *pxdAΔ* hyphae with fluorescently tagged autophagosomes (GFP-Atg8). Scale bar, 5 μm. **(B)** Autophagosome distribution is quantified by fluorescence intensity line-scans of fluorescently-tagged organelles in wild-type or *pxdAΔ* hyphae. Mean fluorescence intensity (solid lines) ± SEM (shading) is plotted as a function of distance from the hyphal tip. n = 23 (WT) and 23 (*pxdAΔ*) hyphae. **(C)** Representative micrographs of wild-type or *pxdAΔ* hyphae with fluorescently tagged mitochondria (Tom20-GFP). Scale bar, 5 μm. **(D)** Mitochondria distribution is quantified by fluorescence intensity line-scans of fluorescently-tagged organelles in wild-type or *pxdAΔ* hyphae. Mean fluorescence intensity (solid lines) ± SEM (shading) is plotted as a function of distance from the hyphal tip. n = 37 (WT) and 31 (*pxdAΔ*) hyphae.

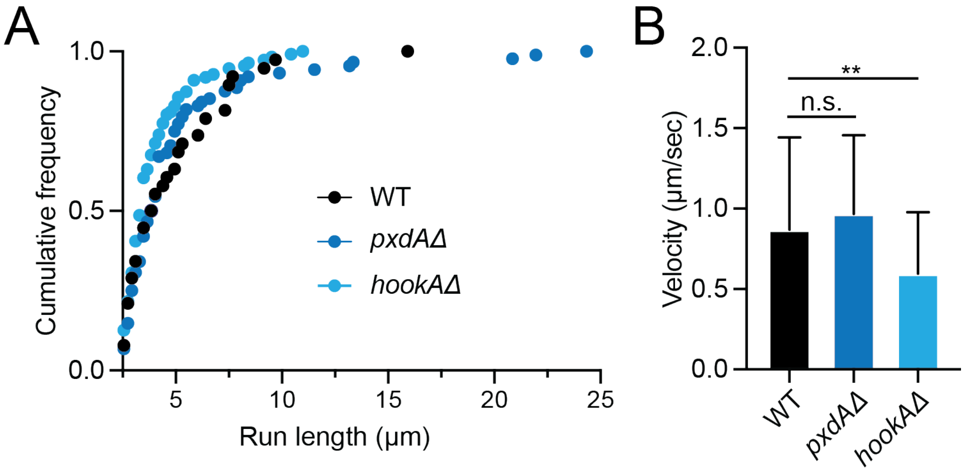


**Supplemental Figure S2: Lipid droplet distribution and movement is not affected by PxdA or HookA.** (**A**) Cumulative distribution of run lengths (x-axis starts at 2.5 µm). n = 38 (WT), 88 (*pxdAΔ*) and 111 (*hookAΔ*) puncta. Distributions are not significant by Kruskall-Wallis test. **(B)** Bar graphs of lipid droplet velocity for puncta displaying processive runs greater than 2.5 µm in wild-type, *pxdAΔ* and *hookAΔ* hyphae. n = 33 (WT), 58 (*pxdAΔ*) and 65 (*hookAΔ*) puncta. WT vs. *hookAΔ* condition is significant (**p = 0.003, one-way ANOVA with Dunnett's posthoc test).

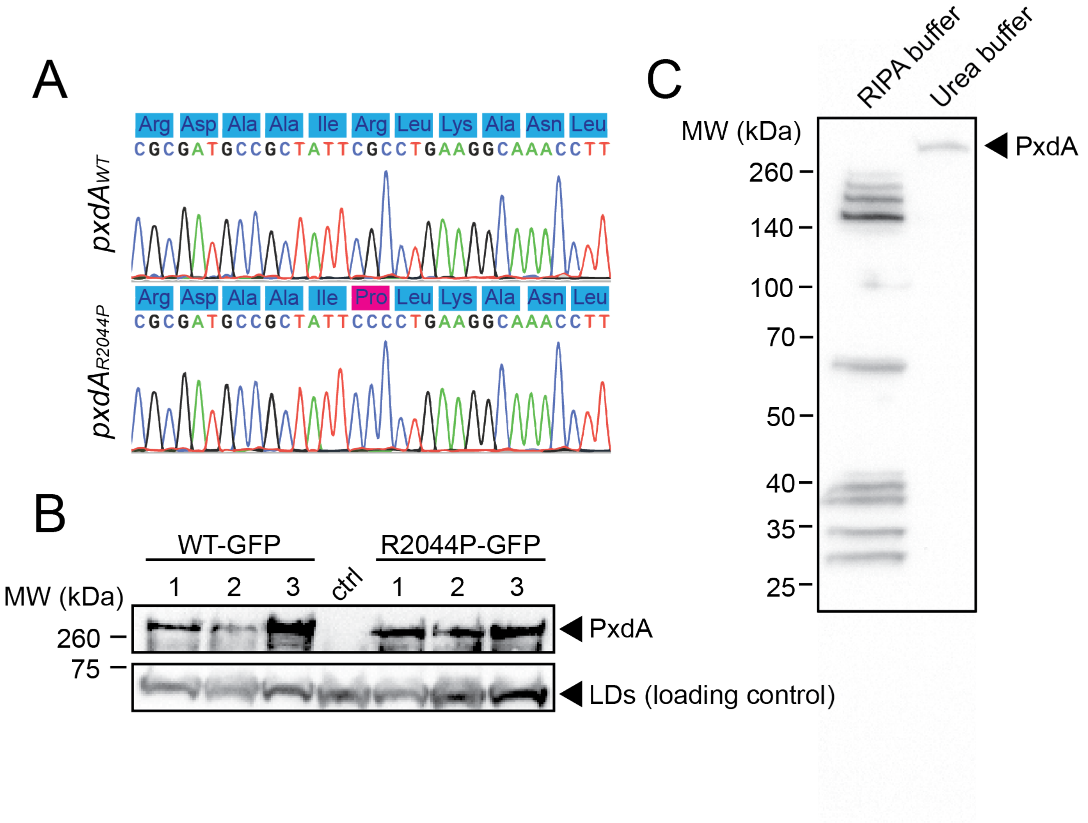


**Supplemental Figure S3: Characterization of PxdA$_{WT}$ and PxdA$_{R2044P}$. (A)** DNA sequence chromatogram and corresponding codons of mutagenized strains from our screen (Tan *et al.*, 2014) displaying a wild-type or point mutation in AN1156/PxdA. Arginine (CGC) at amino acid position 2044 is converted to Proline (CCC). **(B)** Lysates from strains expressing either no tag, PxdA-TagGFP or PxdA$_{R2044P}$-TagGFP were immunblotted for anti-TagGFP. All strains also expressed the lipid droplet marker AN7146-mKate and served as a loading control. Numbers (above Western blot) denote technical replicates for each condition. **(C)** Lysates from strain expressing PxdA-TurboID-3xFLAG resuspended in RIPA or urea buffer immunoblotted with anti-FLAG.

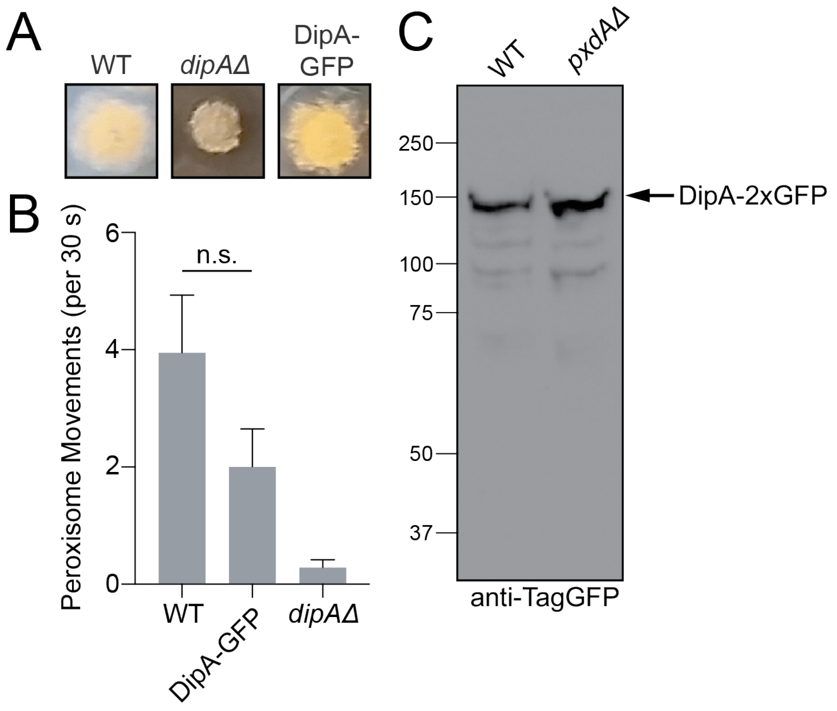


**Supplemental Figure S4: Characterization of *dipAΔ* and DipA-GFP strains. (A)** Colony growth of wild-type, *dipAΔ* and DipA-2xmTagGFP2 (DipA-GFP)-expressing strains. **(B)** Quantification of peroxisome movements (calculated as in Figure 6E) in wild-type, DipA-GFP, and *dipAΔ* strains. Data is mean ± SEM. n = 17 (WT), 9 (DipA-GFP), and 25 (*dipAΔ*). Wild-type and DipA-GFP conditions are not significantly different by Mann-Whitney test. *dipAΔ* data from Figure 6E included for comparison. **(C)** Lysates from wild-type and *dipAΔ* strains expressing DipA-GFP immunoblotted with anti-TagGFP.

**Table S1. *Aspergillus nidulans* strains used in this study.**

All promoters at the endogenous promoter, unless otherwise stated.

| Strain | Genotype | Source |
|---|---|---|
| RPA288 | *yA1*::[*gpdA(p)*-mCherry-FLAG-PTS1::*Afpyro*], *pyroA4, pyrG89, nkuA::Bar* | Salogiannis et al., 2016 |
| RPA520 | *yA1*::[*gpdA(p)*-mcherry-FLAG-PTS1::*Afpyro*], [HH1-BFP::*Afribo*], [TagGFP-RabA*:: AfpyrG*], *pyroA4, pyrG89, pabaA1, nkuA:: argB+* | Tan et al., 2014 |
| RPA568 | [*pxdA$^{R2044P}$*], *yA::*[*gpdA(p)*-mcherry-FLAG-PTS1::*Afpyro*], [HH1-BFP::Afribo], [TagGFP-RabA*:: AfpyrG*], *pyroA4, pyrG89, pabaA1, nkuA:: argB+* | Tan et al., 2014 (mutagenized); Sequenced in this study |
| RPA604 | [*pxdA$^{S201stop}$*], *yA*::[*gpdA(p)*-mcherry-FLAG-PTS1::*Afpyro*], [HH1-TagBFP::*Afribo*], [TagGFP2-*RabA:: AfpyrG*], *pyroA4, pyrG89, pabaA1, nkuA:: argB+* | Tan et al., 2014; Salogiannis et al., 2016 |
| RPA628 | [*pxdA$^{Q846stop}$*], *yA*::[*gpdA(p)*-mcherry-FLAG-PTS1::*Afpyro*], [HH1-TagBFP::*Afribo*], [TagGFP2-RabA:: *AfpyrG*], *pyroA4, pyrG89, pabaA1, nkuA:: argB+* | Tan et al., 2014; Salogiannis et al., 2016 |
| RPA639 | [*pxdA$^{Q1201stop}$*], *yA*::[*gpdA(p)*-mcherry-FLAG-PTS1::*Afpyro*], [HH1-TagBFP::*Afribo*], [TagGFP2-RabA:: *AfpyrG*], *pyroA4, pyrG89, pabaA1, nkuA:: argB+* | Tan et al., 2014 (mutagenized); Sequenced in this study |
| RPA752 | [TagGFP2-AtgH$^{Atg8}$::*Afribo*], *yA1,riboB2, pyroA4, pyrG89, pabaA1, nkuA:: argB+* | This study |
| RPA872 | *yA*::[*gpdA(p)*mcherry-FLAG-PTS1::*Afpyro*], *pyroA4*, [PxdA-TagGFP2::*AfpyrG*], *pyrG89, nkuA::Bar* | Salogiannis et al., 2016 |
| RPA878 | *yA*::[*gpdA(p)*mcherry-FLAG-PTS1::*Afpyro*], *pyroA4*, [PxdA-HA::*AfpyrG*], *pyrG89, nkuA::Bar* | Salogiannis et al., 2016 |

| | | |
|---|---|---|
| RPA915 | [TagGFP2-AtgH$^{Atg8}$::*Afribo*], *yA1,riboB2, pyroA4,* [*pxdAΔ::pyrG*], *pyrG89, pabaA1, nkuA:: argB*+ | This study |
| RPA921 | *yA1,* [*pxdAΔ::Afribo*], *riboB2, yA1*::[*gpdA(p)*-mCherry-FLAG-PTS1::*Afpyro*], *pyroA4, pyrG89, pabaA1, nkuA:: argB*+ | Salogiannis et al., 2016 |
| RPA978 | *yA1*::[*gpdA(p)*-mCherry-FLAG-PTS1::*Afpyro*], *pyroA4,* [PxdA$^{Q2044P}$-TagGFP2::*AfpyrG*], *pyrG89, nkuA::Bar* | This study |
| RPA996 | *yA1,* [TagGFP2-RabA:: *AfpyrG*], *pyrG89,* [PxdA$^{Q2044P}$-mKate::*Afpyro*], *pyroA4, pabaA1, nkuA::argB*+ | This study |
| RPA1002 | *yA1*::[*gpdA(p)*-mcherry-FLAG-PTS1::*Afpyro*], [TagGFP2-RabA::*AfpyrG*], [*dipAΔ*::*Afribo*], *riboB2, pyroA4, pyrG89, pabaA, nkuA:: argB*+ | This study (also see Schinke et al., 2016) |
| RPA1028 | *yA1*::[*gpdA(p)*-mcherry-FLAG-PTS1::*Afpyro*], *pyroA4*, [TagGFP2-PxdA$^{\Delta 1\text{-}500}$::*AfpyrG*], *pyrG89, pabaA1, nkuA::argB or* Bar? | This study |
| RPA1030 | *wA*::[*gpdA(p)*-2xBFP-PTS1::*Afribo*], *riboB2*, [PxdA-mKate2::*Afpyro*], *pyroA4*, [DipA-2xmTagGFP2::*pyrG*], *pyrG89, nkuA::argB*+ | This study |
| RPA1032 | *wA*::[*gpdA(p)*-2xBFP-PTS1::*Afribo*], *riboB2*, [PxdA-mKate2::*Afpyro*], *pyroA4,* [DipA-2xTagGFP2::*AfpyrG*}, *pyrG89*, [*hookAΔ*::Bar], *nkuA:: argB*+ | This study |
| RPA1036 | *yA*::[*gpdA(p)*-mcherry-FLAG-PTS1::*Afpyro*], *pyroA4*, [DipA-2xTagGFP2::*AfPyrG*], *pyrG89*, *nkuA*::Bar | This study |
| RPA1040 | [*dipAΔ*::*Afribo*], [TagGFP2-PxdA$^{\Delta 1\text{-}500}$::*AfpyrG*], *pyroA4, pabaA1, pyrG89, riboB2(?), nkuA::argB* or Bar*?* | This study |
| RPA1045 | *yA1, riboB2, pyroA4, pyrG89, pabaA1*, [DipA-2xmTagGFP2::*AfpyrG*], [*pxdAΔ*::*Afribo*], *nkuA::argB*+ | This study |
| RPA1052 | *yA*::[*gpdA(p)*-mCherry-FLAG-PTS1::*Afpyro*], *pyroA4,* [AN0559$^{Tom20}$-TagGFP2::*AfpyrG*], *pyrG89, nkuA::Bar* | This study |
| RPA1053 | *yA1*,[*pxdAΔ*::*Afribo*], *riboB2, pyroA4,* [AN0559$^{Tom20}$-TagGFP2::*AfpyrG*]*, pyrG89, pabaA1, nkuA:: argB*+ | This study |

| | | |
|---|---|---|
| RPA1212 | yA1, riboB2, pyrG89, [AN7146$^{Erg6}$-mKate::*Afpyro*], pyroA4, pabaA1, nku::argB+ | This study |
| RPA1214 | yA1, riboB2, [PxdA-TagGFP2::AfpyrG], pyrG89, [AN7146$^{Erg6}$-mKate::*Afpyro*], pyroA4, pabaA1, nku::argB+ | This study |
| RPA1215 | yA1, riboB2, [PxdA$^{Q2044P}$-TagGFP2::*AfpyrG*], pyrG89, [AN7146$^{Erg6}$-mKate::*Afpyro*], pyroA4, pabaA1, nku::argB+ | This study |
| RPA1216 | yA1, riboB2, [*pxdAΔ::*AfpyrG], pyrG89, [AN7146$^{Erg6}$-mKate::*Afpyro*], pyroA4, pabaA1, nku::argB+ | This study |
| RPA1218 | yA1, riboB2, [*hookAΔ::*AfpyrG], pyrG89, [AN7146$^{Erg6}$-mKate::*Afpyro*], pyroA4, pabaA1, nku::argB+ | This study |
| RPA1222 | *yA1*::[*gpdA(p)*-mCherry-FLAG-PTS1::*Afpyro*], *pyroA4,* [*hookAΔ::*AfpyrG], *pyrG89, nkuA::Bar* | This study |
