## Supplementary material for "PxdA interacts with the DipA phosphatase to regulate endosomal hitchhiking of peroxisomes": Movie Legends

**SUPPLEMENTAL MOVIE LEGENDS**

Movie S1. **Lipid droplet movements.** Lipid droplet (Erg6/AN7146-mKate) movement in an *A. nidulans* hypha. Arrows denote processively-moving lipid droplets. Images were acquired by time-lapse epifluorescence microscopy using an inverted spinning-disk microscope (Nikon). Frames were taken every 500 ms for 1 min. Video frame rate is 15 frames/s.

Movie S2. **Lipid droplets movement is distinct from early endosome movement**. Early endosome (TagGFP2-RabA) movement, lipid droplet (Erg6/AN7146-mKate) movement, and merged panel (early endosomes – green, lipid droplets – magenta) in an *A. nidulans* hypha. Arrows denote processively-moving lipid droplet. Images were acquired by time-lapse epifluorescence microscopy using an inverted TIRF microscope (Nikon). Scale bar, 5 μm and time in seconds.

Movie S3. **PxdA$_{R2044P}$-expressing hyphae have defects in peroxisome movement.** Peroxisome (mCherry-PTS1) movement in an *A. nidulans* hypha expressing either PxdA$_{WT}$-GFP or PxdA$_{R2044P}$-GFP from the endogenous *pxdA* promoter. Arrows in magenta indicate long-range movement of peroxisomes. Images were acquired by time-lapse epifluorescence microscopy using an inverted TIRF microscope (Nikon). Frames were taken every 500 ms for 45 sec. Video frame rate is 15 frames/s.

Movie S4. **PxdA$_{R2044P}$ foci are less motile and more diffuse.** PxdA-GFP movement from a PxdA$_{WT}$-GFP or PxdA$_{R2044P}$-GFP expressing (endogenous) hypha. Images were acquired by time-lapse epifluorescence microscopy using an inverted TIRF microscope (Nikon). Frames were taken every 367 ms for 20 sec. Video frame rate is 15 frames/s.

Movie S5. **DipA localizes to motile foci.** DipA movement visualized in a hypha expressing DipA-2xTagGFP2 from the endogenous *dipA* promoter. Images were acquired by time-lapse epifluorescence microscopy using an inverted spinning-disk microscope (Nikon). Frames were taken every 400 ms for 20 sec. Video frame rate is 15 frames/s.

Movie S6. **PxdA movements are normal in *dipAΔ* strains.** PxdA (GFP-PxdA) movement in a *dipA* deletion strain. Images were acquired by time-lapse epifluorescence microscopy using an inverted TIRF microscope (Nikon). Frames were taken every 250 ms for 20 sec. Video frame rate is 15 frames/s.

Movie S7. **PxdA is required for proper localization of DipA.** DipA-GFP localization and movement compared in a wild-type and a *pxdA* deletion strain. Images were acquired by time-lapse epifluorescence microscopy using an inverted spinning-disk microscope (Nikon). Frames were taken every 300 ms for 15 sec. Video frame rate is 15 frames/s.

Movie S8. **DipA is required for peroxisome movement.** Visualization of peroxisome movement (arrows in magenta) along a wild-type hypha and a hypha from a *dipA* deletion strain. Images were acquired by time-lapse epifluorescence microscopy using an inverted TIRF microscope (Nikon). Frames were taken every 500 ms for 30 sec. Video frame rate is 15 frames/s.

Movie S9. **DipA is not required for early endosome movement.** Visualization of early endosome movement (GFP-RabA/5a) along a wild-type hypha and a hypha from a *dipA* deletion strain. Images were acquired by time-lapse epifluorescence microscopy using an inverted TIRF microscope (Nikon). Frames were taken every 250 ms for 13 sec. Video frame rate is 15 frames/s.

Movie S10. **DipA cotransports with moving peroxisomes.** Simultaneous two-color time-lapse epifluorescence imaging of a hypha with DipA-GFP (green) and peroxisomes (magenta). Images were acquired using the OMX Blaze v4 (GE Healthcare). Frames were taken every 250 ms for 30 sec. Video frame rate is 15 frames/s.
